# LFCT: A Benchmark Dataset for Low-Frame-Rate Cell Tracking in Long-Term Live-Cell Microscopy

**DOI:** 10.64898/2026.05.30.728955

**Authors:** Mina Gachloo, Tirthankar Biswas, Xiaoming Lu, Caroline M. Greene, Chandler K. Hargett, Katelyn R. Simancik, Marc R. Birtwistle, Federico Iuricich

## Abstract

Cell tracking in time-lapse microscopy is essential for studying dynamic biological processes such as migration, proliferation, and lineage formation. Existing benchmarks primarily focus on high-framerate imaging, where short temporal intervals simplify correspondence between cells across consecutive frames. We present the Low Frame-rate Cell Tracking dataset (LCFT), a benchmark dataset designed specifically for evaluating cell tracking methods under low-frame-rate conditions. The dataset contains multi-day live-cell microscopy sequences from four human cell lines (MCF10A, MDA-MB-231, HEK293T, and U87), acquired at 10× and 20× magnifications using phase-contrast and fluorescence imaging (nucleus). Ground-truth annotations include cell identifications, temporal linking, lineage relationships, and mitosis events. To generate reliable annotations, automated segmentation and tracking were combined with extensive manual curation. LCFT provides a comprehensive resource for developing and benchmarking robust cell tracking algorithms capable of handling sparse temporal sampling and large inter-frame motion in long-term live-cell imaging experiments.

## 1 Background and summary

Automated tracking of individual cells in time-lapse microscopy is a fundamental step for quantitative cell biology, supporting the analysis of proliferation, lineage, migration, cell-cycle dynamics, and response to perturbation [12, 29, 21, 28, 43]. These measurements are essential in common live-cell culture assays such as drug-response screens [37, 5], wound-healing assays [25, 18], stem-cell differentiation studies [33, 4], and co-culture models of cell-cell interaction [19].

Yet, many experiments face a critical tradeoff. Capturing images frequently enough to resolve individual cell events increases phototoxicity [22], a light-induced stress that alters normal cell behavior, accelerates cell death, and compromises the biological validity of the experiment. High imaging frequency also leads to photobleaching [34], the irreversible fading of fluorescent signals due to repeated light exposure, which diminishes image quality and limits long-term observation. Moreover, in high-throughput experiments, high frame rates restrict the number of fields of view or experimental conditions that can be imaged simultaneously, reducing experimental scalability [39].

Lowering imaging frequency could mitigate all these issues, but sparse sampling fundamentally alters the tracking problem. Lower temporal resolution leads to larger cell displacements between frames, reduced spatial overlap, and increased ambiguity in cell correspondence, making automated tracking substantially more challenging [13].

This limitation affects the entire cell-tracking workflow, from automated analysis to human annotation and data exploration. Over the past decade, tracking algorithms have achieved remarkable progress, driven largely by deep learning architectures that have steadily improved the accuracy of cell detection, segmentation, linking, and lineage reconstruction [14, 45, 7, 27, 16, 3]. These advances have enabled increasingly reliable analysis of large-scale microscopy datasets and have become the foundation of modern cell-tracking pipelines.

In parallel, a rich ecosystem of visualization, annotation, and analysis software has emerged to support researchers in managing and curating tracking data [36, 38, 23, 15, 31, 41, 1, 26]. These tools provide interactive interfaces for reviewing trajectories, correcting tracking errors, and exploring lineage relationships, making them essential components of the dataset generation and validation process.

However, both automated tracking methods and user-facing analysis tools have been developed primarily for densely sampled microscopy sequences, where cells undergo relatively small displacements between consecutive frames. Their underlying assumptions, algorithms, and interaction paradigms are therefore optimized for high temporal resolution data. When applied to sparse acquisitions, where cells may move substantial distances between observations and mitosis events can occur between frames, these assumptions no longer hold [13]. As a result, tracking performance deteriorates, lineage reconstruction becomes more challenging, and manual curation interfaces provide less effective support for resolving ambiguities.

In this work, we introduce a benchmark dataset for cell tracking under low-frame-rate imaging conditions. While existing datasets [9, 30] and evaluation frameworks [44] have largely focused on densely sampled microscopy sequences, there is currently no standard resource for assessing performance when temporal resolution is substantially reduced. By combining manually curated ground-truth annotations with a systematic sparsification framework, our dataset enables objective benchmarking across a range of low-frame-rate scenarios. We anticipate that this resource will serve as a stepping stone toward the development of a new generation of tracking algorithms, annotation interfaces, and analysis tools designed explicitly for sparse temporal sampling and long-term live-cell imaging experiments.

The dataset consists of multi-day time-lapse microscopy sequences with ground-truth annotations for cell identification [11], cell linking [2], and mitosis events [24, 46]. To enable evaluation across varying levels of temporal sparsity, we provide a sparsification framework that systematically reduces frame rate while preserving annotation accuracy. Together, these resources establish an objective foundation for developing and evaluating tracking methods capable of handling the large cell displacements and limited temporal context characteristic of sparsely sampled microscopy data.

## 2 Method

The objective of this work is to construct a benchmark dataset for cell tracking under low-framerate imaging conditions. Direct analysis of low-frame-rate microscopy data is particularly challenging because large temporal gaps between consecutive frames increase uncertainty in cell correspondence and can result in incomplete or inaccurate lineage annotations. To address this challenge, we acquired live-cell microscopy sequences over multiple days at a sampling interval of 15 minutes. This data sampling regime allowed us to generate accurate annotation of cell trajectories and cell events, providing reliable ground-truth lineage information. An overview of the dataset generation process is shown in Figure 1. Successively, densely sampled sequences were systematically sparsified by removing frames at regu-lar intervals to emulate progressively lower imaging frequencies. The corresponding lineage annotations were sparsified in the same manner, ensuring consistency between the image data and the ground-truth labels while preserving accurate cell correspondences. The resulting dataset contains multiple low-frame-rate versions of each original sequence while preserving the corresponding ground-truth trajectories, enabling objective benchmarking of cell-tracking methods.

**Figure 1:**
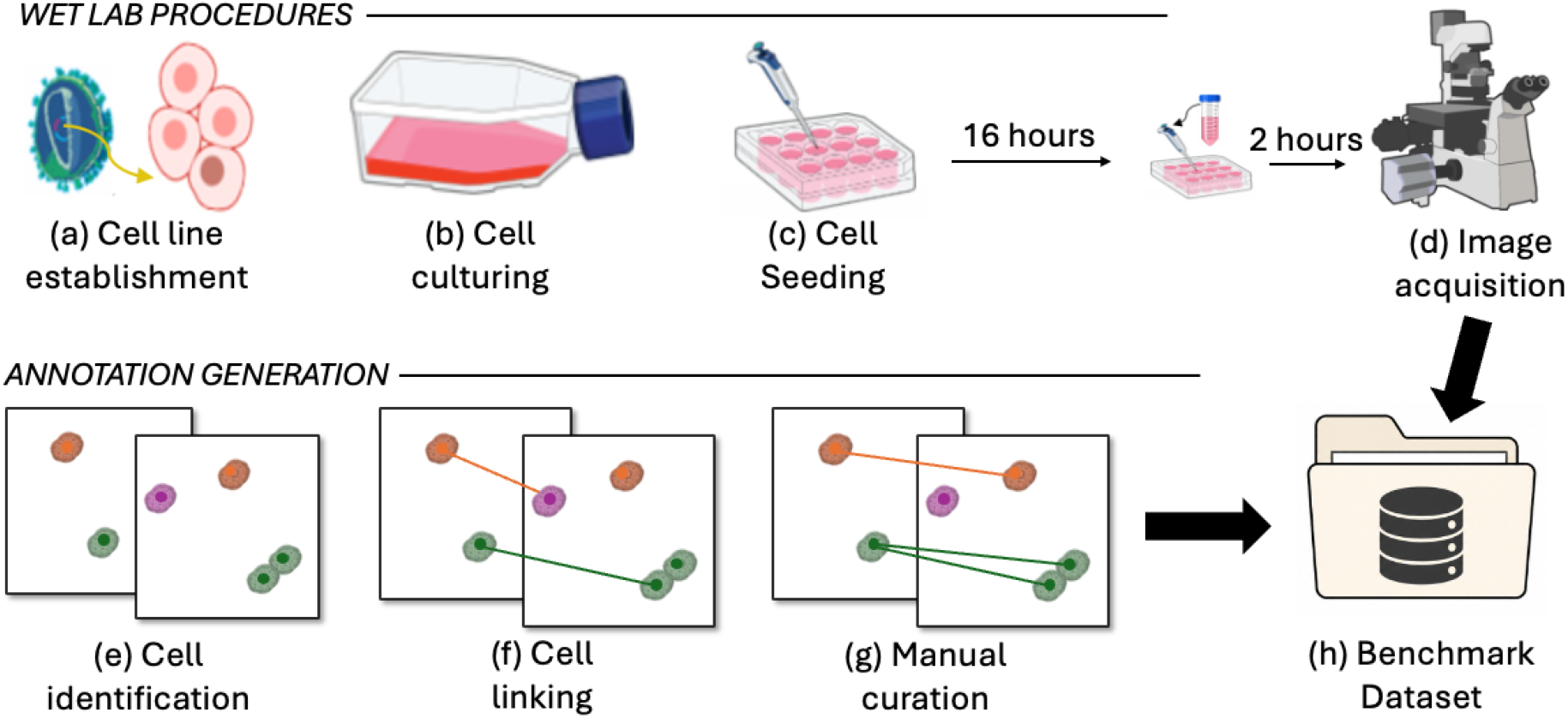
Dataset generation pipeline. Wet-lab Procedures: (a) The H2B-mRuby fluorescent reporter was transduced into each cell line for nuclear labeling. (b) Each line was expanded in growth medium. (c) Cells were seeded into multiwell plates, and the medium was refreshed before imaging. (d) Phase-contrast and fluorescence sequences were acquired at 10× and 20× magnification over three to four days. *Annotation and Benchmark Generation*: (e) Cellpose segmented nuclei and whole cells from the two channels. (f) Detections were linked across frames with the LAP tracker in TrackMate. (g) Tracks were manually curated in TrackScheme to correct linking and identification errors. (h) Images and annotated tracks collected across four cell lines are provided in a new benchmark dataset.

The following subsections describe the experimental and computational pipeline used to generate the dataset. First, we present the procedures for cell culture, fluorescent labeling, and long-term microscopy acquisition 2.1. We then describe the annotation pipeline used to generate ground-truth annotations 2.2. Finally, we describe the sparsification approach we used to simulate sparse acquisition regimes 2.3.

### 2.1 Cell Culture and Imaging Procedures

This dataset contains four human cell lines:

- MCF10A, a non-transformed epithelial cell line;
- MDA-MB-231, a triple-negative breast cancer cell line;
- HEK293T, a human embryonic kidney cell line;
- U87, a human glioblastoma cell line.

#### Cell line establishment and sample preparation

To enable fluorescence-based nuclear tracking, we generated stable H2B-mRuby expressing cell lines via lentiviral transduction.

##### Lentivirus production

A lentiviral vector was used to deliver the pLenti-puro-H2B-mRuby plasmid. For third-generation lentivirus production [8], 5 × 10^6^ HEK293T cells were seeded into a T75 flask (Greiner #658175) and incubated for 22 hours prior to transfection. Media was replaced with serum-free DMEM two hours before co-transfection, which was performed using 6000 ng of pPAX (Addgene #12260), 3000 ng of pCMV-VSV-G (Addgene #8454), and 15000 ng of transfer plasmid with Lipofectamine 2000 (Thermo-Fisher #11668027) per the manufacturer’s protocol. Full growth media (DMEM, 10% FBS, 2 mM L-glutamine) was restored 6 hours post-transfection. Lentiviral supernatant was collected at 48 and 72 hours post-transfection, pooled, and centrifuged at 1000 × *g* for 5 minutes to remove cell debris. The supernatant was concentrated using an Amicon Ultra-15 100 kDa centrifugal filter (Millipore #UFC910008), aliquoted, and stored at −80°C [6].

##### Lentiviral transduction

A suspension of 1 × 10^5^ cells was transduced with 150 µL of concentrated lentiviral supernatant in full growth media in a 6-well plate (Corning Falcon #353046). Fluorescence expression was verified by imaging 48 hours post-transduction. All cell lines were selected with puromycin (2 µg/mL, STEMCELL #73342) for one week until non-transduced control cells were dead. Stable H2B-mRuby expressing cells were subsequently maintained in full growth medium supplemented with 0.2 µg/mL puromycin [40].

#### Image acquisition

All images were acquired using a BioTek Cytation 5 under environmental control conditions (37 °C and 5% CO_2_). All cell lines expressed H2B-mRuby as a nuclear marker. During the imaging intervals, cells were maintained in a BioTek BioSpa 8 automated incubator under the same environmental conditions (37 °C and 5% CO_2_).

For imaging, all cell lines were seeded into 12-well plates (Corning Falcon, #353043). The seeding density for HEK293T cells was 3–4 × 10^4^ cells/well, whereas MCF10A, MDA-MB-231, and U87 cells were seeded at 1.5–3 × 10^4^ cells/well. Cells were incubated for 16 hours at 37 °C and 5% CO_2_, after which the medium was replaced to remove floating cells, followed by an additional 2-hour incubation. Images were acquired at 10× and 20× magnifications using the RFP channel (EX 531/40 nm, EM 593/40 nm) to visualize mRuby fluorescence, along with the phase-contrast channel, at 10–15 minute intervals over three to four days.

For each cell line, we used two imaging conditions: healthy and unhealthy. The unhealthy condition applied 100% LED intensity with varying integration time, to ensure lack of proliferation and in some cases extensive death. The healthy condition applied 30% LED intensity for U87 and 10% LED intensity for MDA-MB-231 and HEK293T, also with varying integration time. The unhealthy condition was applied to the 10× sequences of MCF10A, HEK293T, and MDA-MB-231, together with two MDA-MB-231 sequences at 20×, while all remaining sequences were imaged under the healthy condition. Table 1 reports the number of annotated sequences for each cell line and magnification.

**Table 1:** Number of annotated sequences per cell line and magnification.

| Cell line | $10\times$ | $20\times$ | Total |
| --- | --- | --- | --- |
| MCF10A | 9 | 9 | 18 |
| HEK293T | 8 | 6 | 14 |
| MDA-MB-231 | 7 | 6 | 13 |
| U87 | 4 | 4 | 8 |

### 2.2 Annotation and Benchmark Generation

To generate reliable ground-truth annotations, we employed a semi-automated pipeline integrating automated methods and manual curation at every stage of the annotation process, including cell identification, cell linking, and lineage reconstruction.

#### 2.2.1 Cell identification

##### Initial segmentation

All images were processed using Cellpose [41], a deep learning–based framework for automated cell segmentation [32]. The two acquired imaging channels, fluorescence and phase-contrast, were processed independently using pre-trained Cellpose models specialized for different cellular structures. Specifically, the *nuclei* model was applied to the fluorescence images to segment nuclear regions, while the *Cytoplasm (cyto2)* model [35] was used on the phase-contrast images to delineate whole-cell boundaries. Models were downloaded from the official Cellpose repository (https://github.com/MouseLand/cellpose), which provides them through the Cellpose python package. Figure 2(a-d) illustrates representative outputs from both models, showing the fluorescence and phase-contrast inputs alongside their corresponding segmentation masks. Each frame was independently processed to handle changes in cell density and morphology over time.

**Figure 2:**
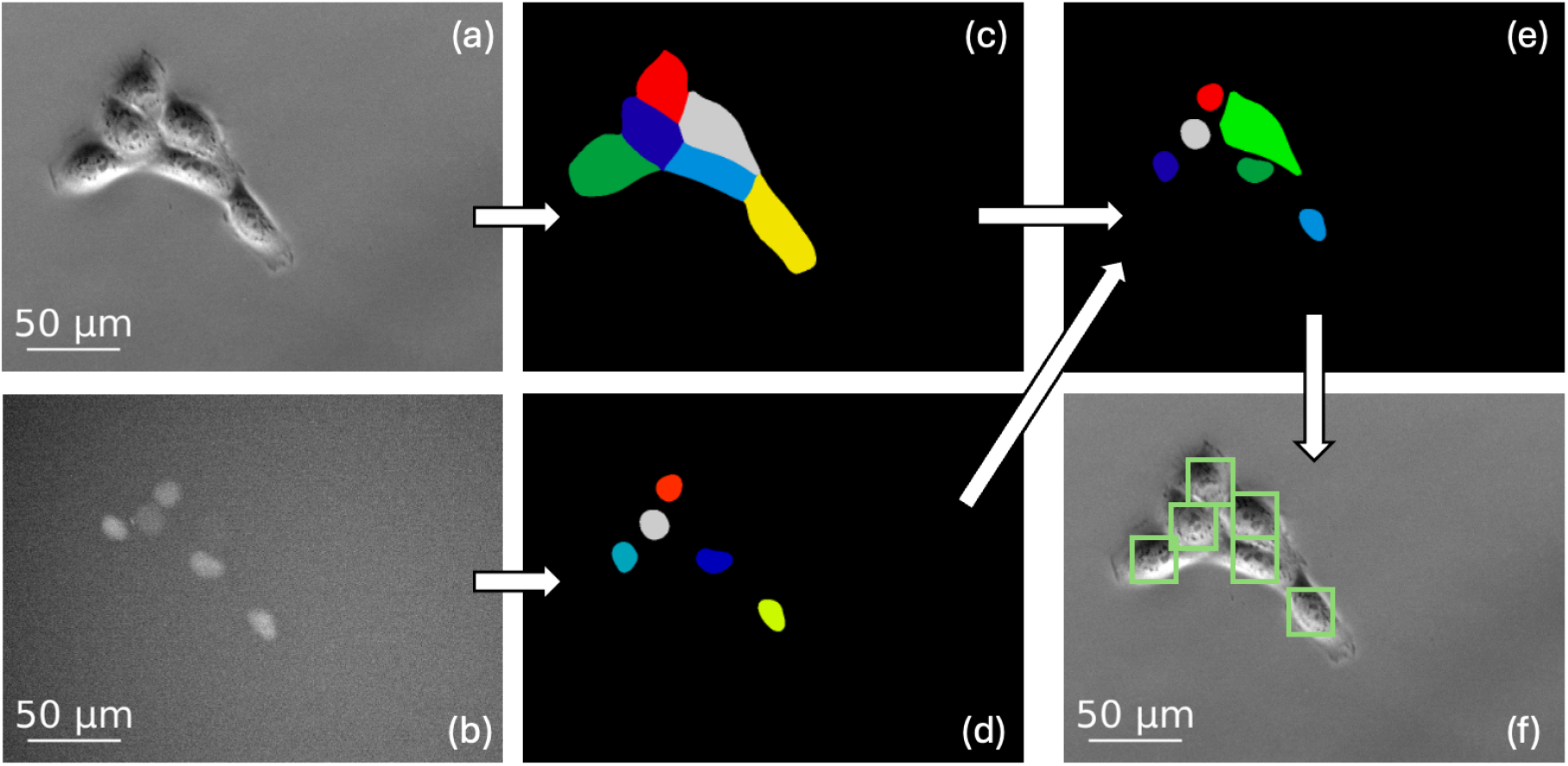
Example of the cell identification pipeline applied to a cropped region from an MCF10A sequence acquired at 20× magnification. The fluorescence and phase-contrast channels were processed independently using Cellpose and subsequently integrated to obtain a single cell representation. (a) Raw phase-contrast image. (b) Raw fluorescence image showing H2B-mRuby–labeled nuclei. (c) Whole-cell segmentation generated by the Cellpose cyto2 model. (d) Nuclear segmentation generated by the Cellpose nuclei model. (e) Integrated segmentation obtained by combining the outputs of both models, with one region assigned to each detected cell. (f) Cell locations extracted from the integrated segmentation and used for downstream tracking and lineage annotation.

##### Mask integration

Following the initial segmentation step, we integrated the nuclear and cytoplasmic masks to generate a unified representation of cell position. Because the two Cellpose models were applied independently, each model occasionally failed to detect cells in specific frames due to variations in morphology, cell density, image contrast, or fluorescence signal. We leveraged the complementary detections produced by each model to improve the localization accuracy and reduce manual corrections. To achieve this, we used a hierarchical integration approach. For each detected nucleus, we identified the overlapping cytoplasmic region by determining which cytoplasm mask had the greatest spatial overlap with that nucleus. Figure 2(e) shows an example of this integration, illustrating the combined masks and their boundaries. When a nucleus was successfully matched to a cytoplasmic region, the nuclear boundary was used as the primary cell identifier, as it provides a more consistent and well-defined marker for tracking [17]. In cases where a cytoplasmic region had no associated nucleus, the full cytoplasmic boundary was retained as the cell identifier.

Following mask integration, we imported the resulting masks into Fiji TrackMate [38, 42], an open platform for single-particle tracking, where the Mask Detector [10] converted each labeled region into a detection object (see Figure 2(f)). In case some cell were still missed by the automated segmentation we reviewed each sequence frame by frame and added them as new spots at the corresponding positions, shown in. The complete set of detections was saved as a TrackMate XML file, which used for cell linking.

#### 2.2.2 Cell linking

After registering a location for each cell, we performed cell linking using TrackMate. We employed the Linear Assignment Problem (LAP) tracker [20] to establish correspondences between cells across consecutive frames. We manually fine-tuned two key parameters to optimize the linking results. The frame-to-frame linking maximum distance was adjusted to control the maximum distance a cell can move between frames. The track segment splitting maximum distance was set to identify mitosis events, where a parent cell divides into two daughter cells. This automated linking step produced initial cell trajectories, which required further refinement to correct errors and ensure data quality.

##### Manual Curation

With the preliminary tracking data available, we used the TrackScheme interface to manually review and refine the cell tracks [36]. Modifications within TrackScheme involved adding or deleting tracks and individual cell identifications. During this curation process, we identified and resolved errors in both cell identification and cell linking (Figure 3).

**Figure 3:**
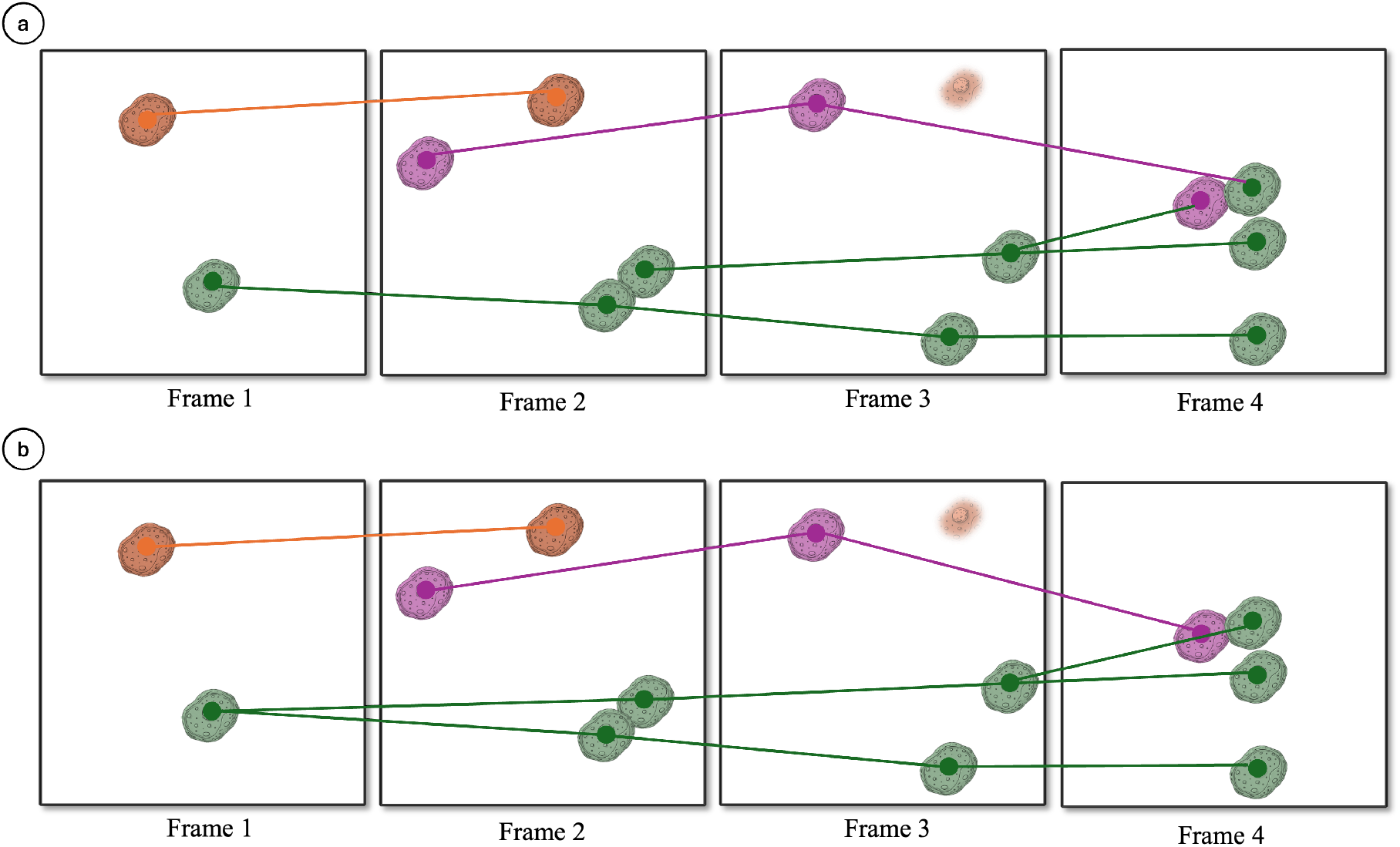
Linking errors and their corrections through manual curation. (a) Three error types from the automated linker: the purple cell links to a wrong cell in frame 4 (false link); the green cell divides in frame 2 but the split is not recorded (false mitosis). (b) The corrected sequence. The purple cell is correctly linked across all four frames. Both daughters of the green cell are linked to the parent after division.

*False links* occurred when a cell was incorrectly associated with a nearby but unrelated cell due to large inter-frame displacement or spatial proximity. We corrected these errors by deleting the incorrect links, reassigning the correct associations, and ensuring each cell maintained a consistent trajectory throughout the sequence.

*False mitosis* happened when the tracker failed to detect a cell in certain frames, breaking the lineage and treating the subsequent detection as a new track. To restore continuity, we manually inserted the missing detections and reconnected the fragmented tracks to their original lineage.

Once the manual curation was complete, we exported the finalized tracks as XML files.

### 2.3 Low-frame-rate generation

With the manually curated ground-truth annotations in place, we can systematically generate lowframe-rate versions of the dataset while preserving accurate cell identities, trajectories, lineage relationships, and mitosis events. Starting from the densely sampled microscopy sequences, we simulate progressively lower temporal resolutions by sparsifying the image sequences and updating the corresponding annotations accordingly. This enables controlled evaluation of cell tracking methods under varying levels of temporal sparsity, providing an objective benchmark for accuracy evaluation.

#### Gap factor

To generate ground-truth data for low-frame-rate conditions, we developed a sparsification approach that systematically downsamples high-frame-rate sequences while preserving accurate cell tracking annotations.

The level of sparsification is defined by the *gap* factor which indicates the number of frames to drop. Specifically, gap x indicates that only one out of every x frames is retained in the sequence. For example, a gap factor of 8 indicate only 1/8 images are retained. Applied to images captured at 15-minute intervals this would result in a temporal resolution of 2 hours between frames.

To maintain consistency between sparsified images and their corresponding annotations, we adjusted the cell tracking annotations to reflect the removed frames. We represented these annotations as lineage trees. Each tree describes the lineage of a cell and its progeny, with each node corresponding to a single cell identification in a given frame and each edge representing either a link between the same cell across consecutive frames or a parent–daughter relationship at a mitosis event. Within a tree, the root node marks the first frame in which the cell appears, internal nodes with multiple children correspond to mitosis events, and leaf nodes indicate cell death or exit from the field of view.

When a frame is removed during sparsification, every cell identification in that frame is also removed from its lineage tree. Each affected tree is then adjusted according to one of three rules, depending on the type of node removed. If the removed node is an internal node, its child nodes are reconnected to its parent node, preserving the overall lineage structure (see the green cell in Figure 4). If the removed node is a root, its child nodes become new root nodes, potentially splitting the lineage into multiple trees (see the purple cell in Figure 4). If the removed node is a leaf, it is simply removed from the tree with no further modifications (see the orange cell in Figure 4).

**Figure 4:**
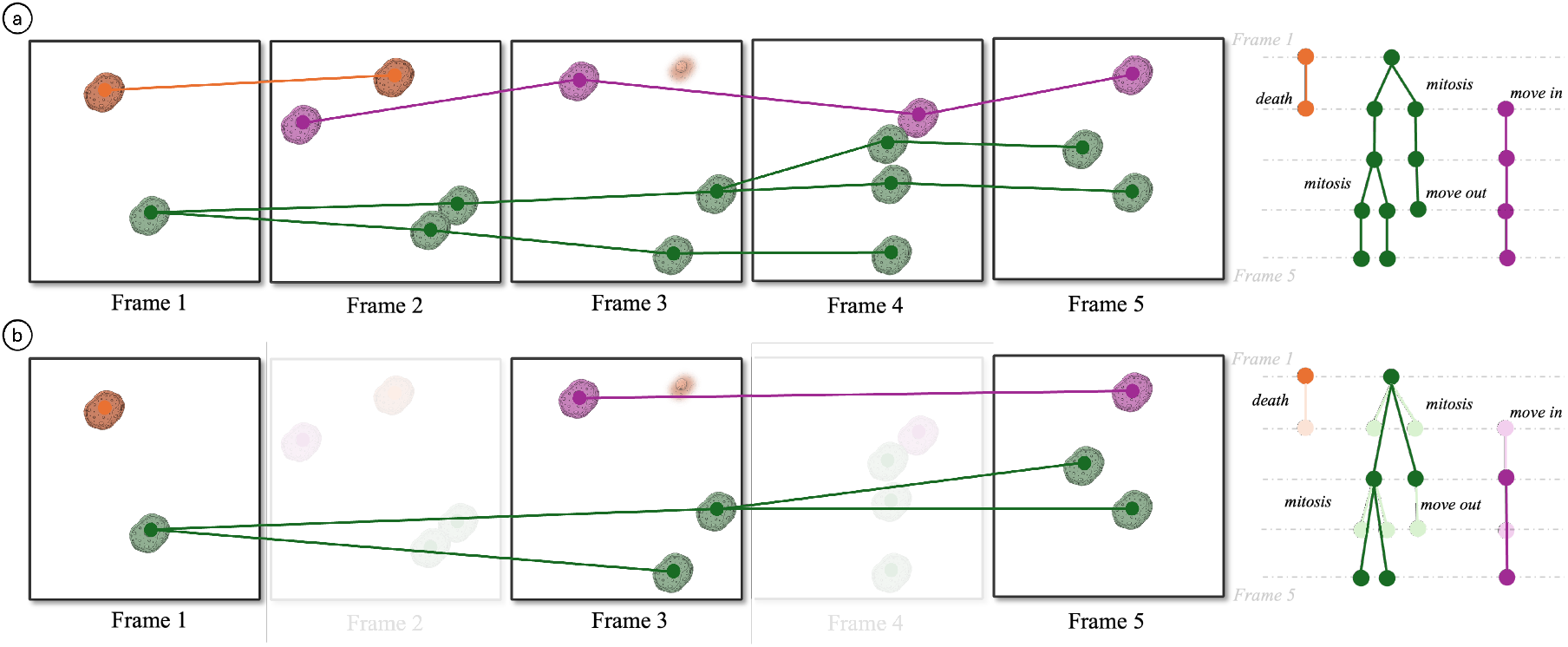
An example of frame sparsification on five frames from a high-frame-rate sequence, using a gap factor of 2. (a) All frames are present, and each cell is linked to its detection in the next frame. The corresponding lineage trees are shown on the right. (b) Every second frame is removed (shown faded) to simulate low-frame-rate conditions, and the lineage trees are adjusted accordingly. Faded nodes correspond to cells in the removed frames.

## 3 Data Availability

The LFCT dataset is available on Zenodo at https://doi.org/10.5281/zenodo.20186357 under the CC-BY 4.0 license. We provide one ZIP file per cell line. Each ZIP file contains:

1. Phase-contrast and nuclear fluorescence time-lapse images in TIFF format.
2. Ground-truth annotations in TrackMate XML format.
3. Time-lapse videos at four GAP factors (1, 2, 4, and 8).

The dataset is also accessible through the project website at https://davislab.github.io/LFCT-dataset. The website is organized by cell line, with one page per cell line that lists all corresponding sequences. For every sequence, the phase-contrast images, fluorescence images, annotations, and video sample can be downloaded individually.

## 4 Technical Validation

This section presents the technical validation of the dataset, with the goal of assessing both the quality of the ground-truth annotations and their biological plausibility. Specifically, we sought to verify that the annotated cell trajectories, lineage relationships, mitosis events, and population dynamics are consistent with the expected behavior of the underlying cell lines.

Section 4.1 presents the validation results for the primary benchmark dataset, which consists of healthy cell populations. In addition, the dataset includes sequences acquired under unhealthy conditions, providing examples of altered cellular behavior. The characteristics of these sequences and their associated annotations are described separately in Section 4.2.

Beyond the quantitative analyses reported in this section, every sequence was subjected to visual inspection. Microscopy videos were reviewed with cell detections, trajectories, and lineage annotations overlaid to verify temporal consistency and identify potential annotation errors. To facilitate independent assessment by dataset users, these annotated videos are provided through the dataset website (see Data Availability), allowing researchers to preview the sequences and corresponding annotations before downloading the data.

### 4.1 Validation of Tracking Dynamics and cell movements

To assess the biological consistency of the annotations, we examine both track characteristics, population growth, and cell movement dynamics.

#### Track and lineage characteristics

The following paragraphs examine the temporal characteristics of the annotated trajectories throughout the imaging sequences.

##### Track lifespan

For each ground-truth track, we analyzed the track onset, termination, and duration within the acquisition sequence. Figure 5 represents each track as a horizontal line spanning the frames in which the corresponding cell is present. Tracks are grouped according to their temporal relationship with the sequence boundaries. The first group (dark green) corresponds to cells that are present throughout the entire sequence, appearing in the first frame and remaining visible until the last frame. The second group (light green) contains tracks that terminate in the final frame but enter the field of view after the sequence begins. The third group (yellow) represents tracks that are already present in the first frame but disappear before the end of the acquisition. Finally, the fourth group (red) includes cells that both enter and exit during the sequence, meaning they are absent from both the first and last frames.

**Figure 5:**
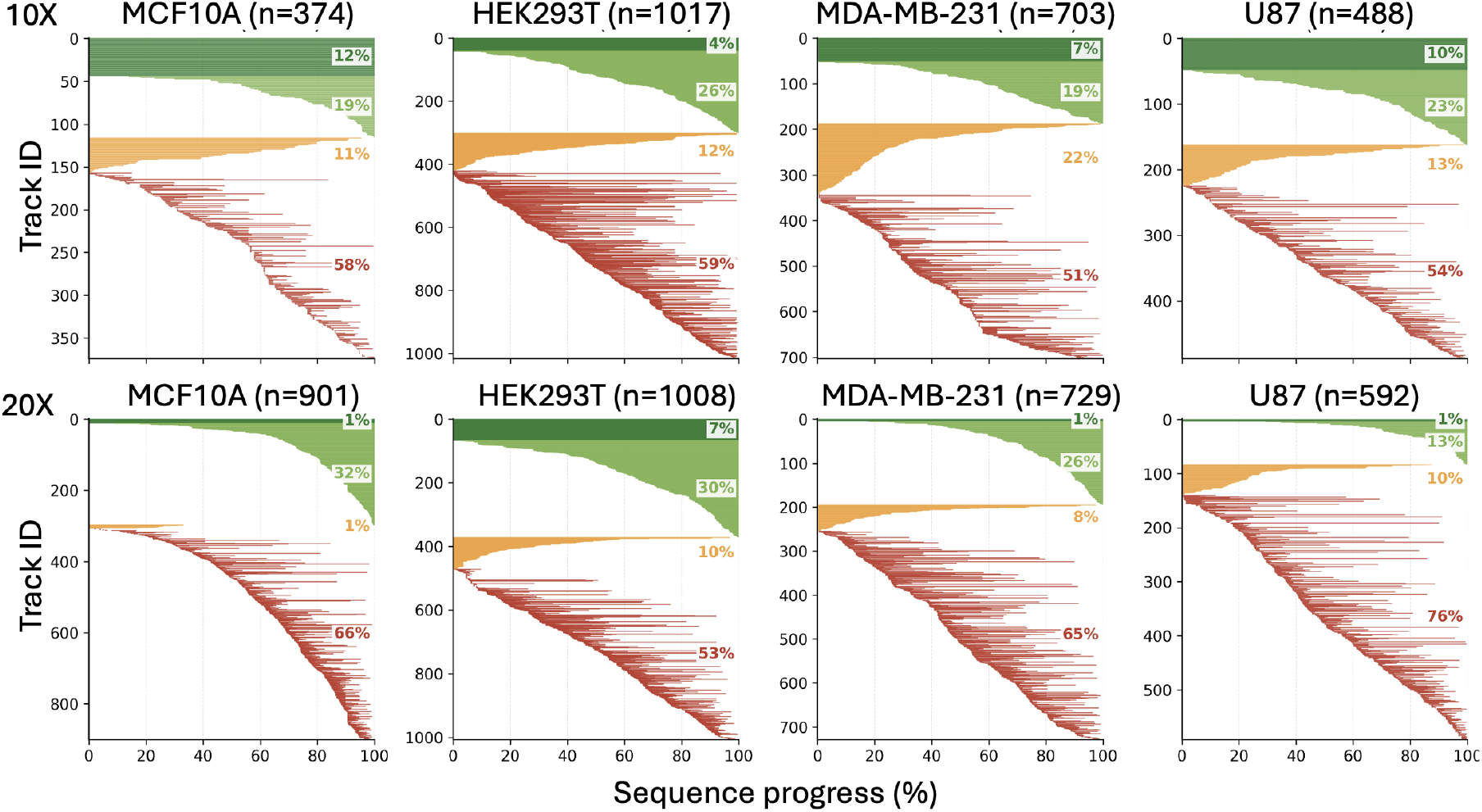
Track presence across the four cell lines imaged at 10× (top row) and 20× (bottom row) magnification. Each horizontal line represents a single cell track and spans the frames in which the cell is present. Tracks are grouped into four categories: cells continuously present throughout the acquisition (dark green), entering after the first frame and remaining until the end (light green), present from the first frame but exiting before the end (orange), and both entering and exiting during the sequence (red). Percentages indicate the proportion of tracks belonging to each category. Each subfigure is labeled with the corresponding cell line and the total number of annotated tracks (*n*)

The four categories together capture both the stability of the cell population in each sequence and how often cells enter and leave the field of view. Across all panels, most tracks belong to cells that enter and exit during the sequence, accounting for 51% to 59% at 10× and 53% to 76% at 20×. The observed distribution of track categories is consistent with expected microscopy acquisition behavior. Cells that remain visible for the entire duration of the sequence are comparatively rare, representing only 4% to 12% of tracks at 10× magnification and 1% to 7% at 20×. The reduced proportion at higher magnification is expected because the smaller field of view at 20× increases the likelihood that cells enter or leave the imaging region during the acquisition.

Figure 6 shows the distribution of track durations for each cell line, where track duration is defined as the number of frames between a cell’s first and last appearance in the sequence. Most tracks are short, with median durations of 40-60 frames at 10× and 20-40 frames at 20×. However, each cell line also includes much longer tracks, with some trajectories extending beyond 400 frames. MDA-MB-231 has the longest typical durations in both panels, and MCF10A contains the longest individual tracks at 10×. The shorter durations at 20× reflect the smaller field of view at higher magnification.

**Figure 6:**
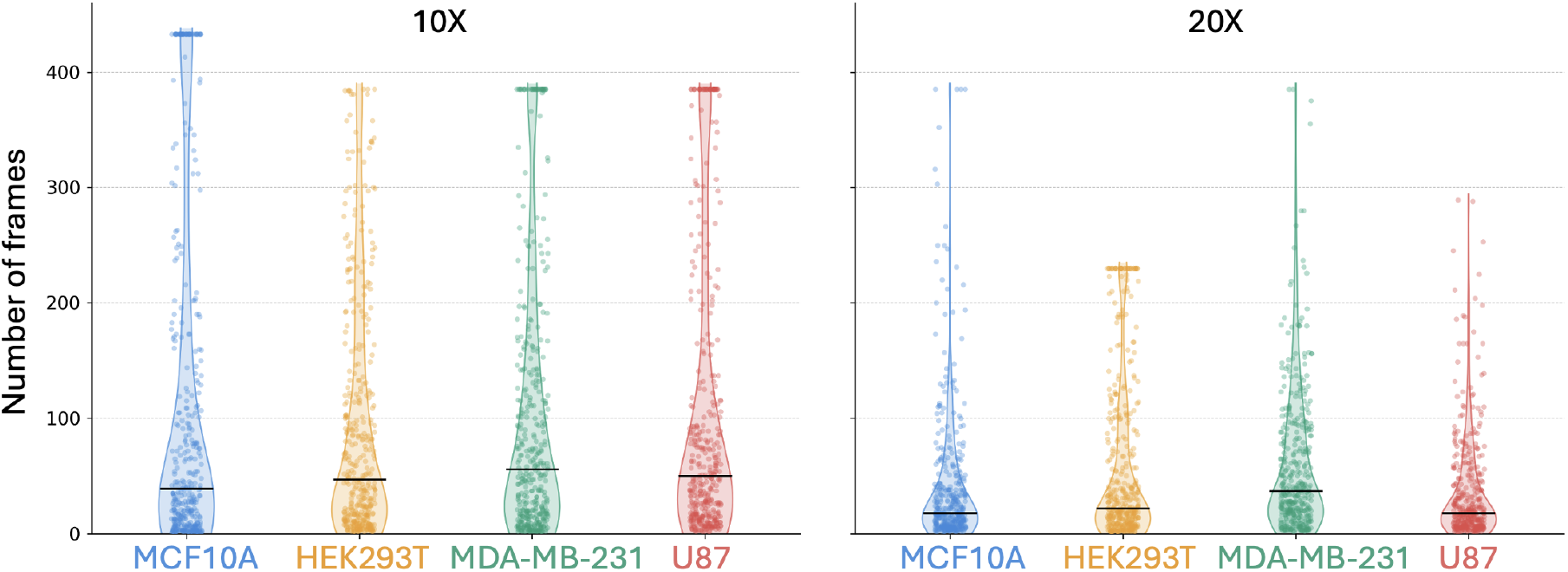
Track duration distribution for the four cell lines at 10× (left) and 20× (right) magnification. Each violin shows the shape of the duration distribution, individual dots show the duration of each track, and the black horizontal line marks the median.

Overall, these results confirm that the annotated trajectories capture realistic population dynamics. The dataset includes a wide range of track durations, from short observations to long trajectories, which is important for evaluating cell tracking methods across different track lengths.

In the healthy condition, cells are imaged under standard culture conditions, stay attached to the substrate, divide regularly, and form a growing population over time. The annotations in this dataset record each cell division and follow every cell across all healthy sequences. The following sections report how often cells divide in each cell line and how the cell population grows over the course of the acquisition.

##### Population growth

The four cell lines also differ in how quickly their populations grow. We measure this by counting the number of cells per unit area at every frame, and report the average across all healthy sequences in Figure 7(a). HEK293T exhibits the highest proliferation rate among the four cell lines and reaches the greatest cell density over the course of the experiment. On average, the number of cells increases from approximately 28 at the beginning of a sequence to more than 115 by its end.

**Figure 7:**
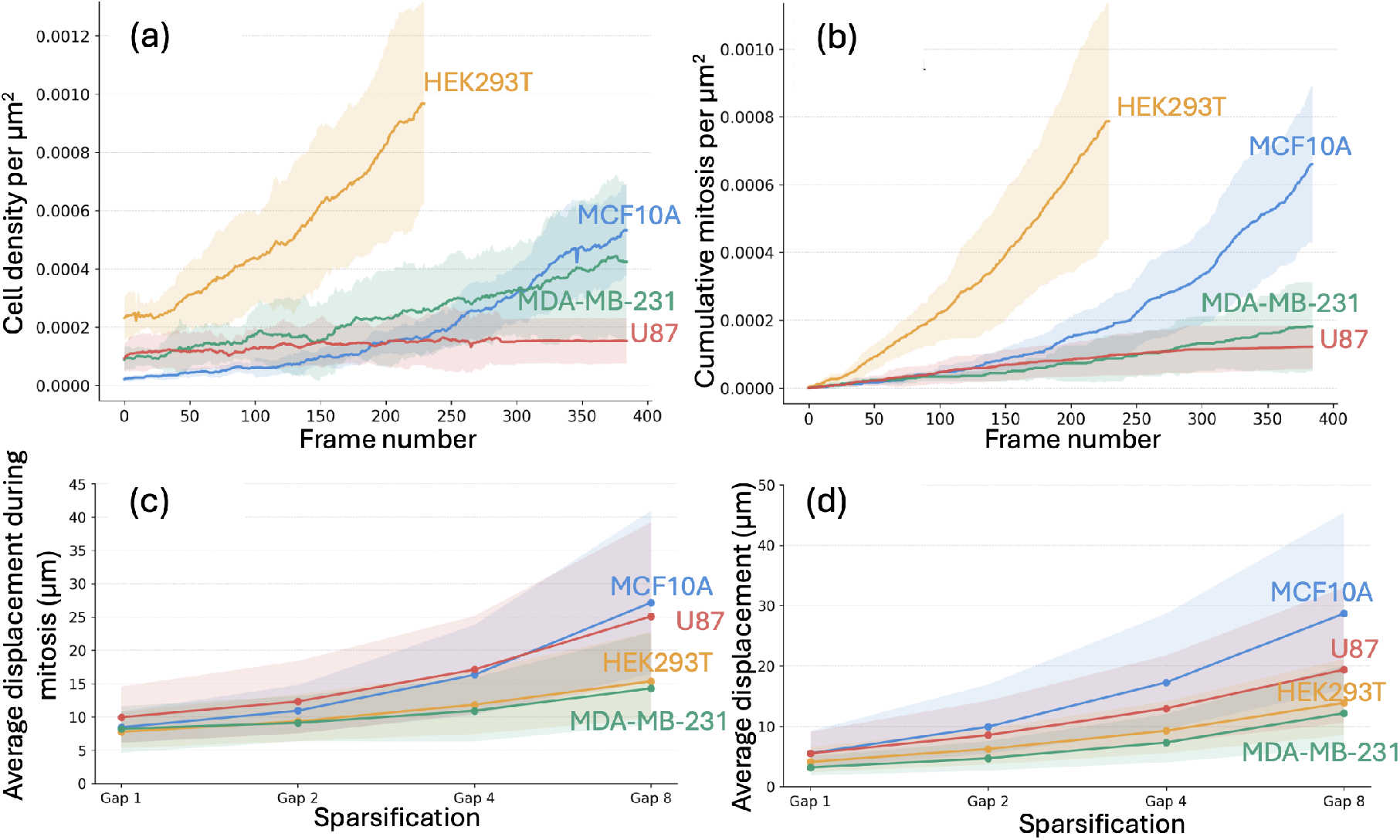
Overview of the healthy cell-line benchmark. (a) Average cell density per unit area and (b) cumulative mitotic events per unit area measured throughout the acquisition period for each cell line. (c) Average parent-to-daughter displacement after mitosis and (d) average displacement between consecutive observations at increasing gap factors. Shaded regions represent variability across sequences of the same cell line.

MCF10A also shows substantial population growth, increasing from roughly 3 cells at the start of a sequence to approximately 65 cells by the final frame. In contrast, MDA-MB-231 displays more moderate growth, with the average cell count increasing from about 11 to 50 cells. U87 exhibits the slowest population expansion, maintaining a relatively stable population of around 22 cells throughout the imaging period.

##### Mitosis progression

Figure 7(b) reports the cumulative number of mitotic events per unit area for each cell line throughout the acquisition. All four cell lines exhibit a steady increase in the number of recorded divisions over time. HEK293T displays the highest mitotic activity, reaching approximately 94 cumulative divisions on average by the end of the acquisition. MCF10A follows closely with roughly 79 divisions, whereas MDA-MB-231 and U87 exhibit substantially lower mitotic activity, accumulating approximately 22 and 32 divisions, respectively.

These differences are consistent with the proliferation dynamics observed in the cell density analysis and reflect the distinct growth characteristics of the four cell lines. Interestingly, U87 and MDA-MB-231 exhibit similar levels of mitotic activity, yet MDA-MB-231 reaches substantially higher cell densities. One possible explanation is that U87 cells are more motile, causing a larger fraction of the population to move out of the field of view during the acquisition. This hypothesis is consistent with the displacement measurements reported in the following section, where U87 displays greater cell movement than MDA-MB-231.

##### Frame-to-frame displacement

To quantify how cells migrate across the field of view in the healthy condition, we compute the displacement of each cell to its position in the next frame and report the distribution per cell line in micrometers. To distinguish migration from division, we exclude the first frame of each daughter cell, since its position is shifted from the parent at the moment of division.

MCF10A and U87 confirms to be the most motile cells with an average frame-to-frame displacement of approximately 7.3 µm each. The average displacement for HEK293T is more moderate 5.0 µm while MDA-MB-231 is the less motile cell cell line in our acquisitions with only 4.0 µm frame-to-frame displacement on average.

These results confirm that the dataset captures a broad range of cell displacement magnitudes, consistent with the trends observed in the cell density and mitosis analyses. They also provide insight into the relative difficulty of the tracking problem across sequences, identifying conditions that are expected to become particularly challenging as temporal sampling is reduced.

##### Sparsification and Low-frame-rate Benchmark

Low-frame-rate imaging reduces the number of observations available per cell and increases cell displacement between consecutive visible frames. Larger displacements make cell linking harder, and fewer frames mean that fewer mitosis events are captured. To show that our dataset captures these conditions, we measured cell displacement between consecutive visible frames and the fraction of mitosis events retained in the sparsified sequences. All these information are reported across gap factors of 1, 2, 4, and 8, for all four cell lines. We recall that gap x indicates that only one out of every x frames is retained in the sequence (Gap 1 is the original sequence).

Figure 7(c) reports the displacement between a parent cell and its daughter cells at the first observable frame following a mitotic event. As the gap factor increases, this displacement grows substantially because the daughter cells continue to migrate during the longer interval between consecutive observations. Consequently, the measured distance reflects both the spatial separation introduced by cell division and the subsequent movement of the daughter cells.

At a gap factor of 8, these distances become particularly large, highlighting the increased difficulty of associating daughter cells with their parent under low-frame-rate conditions. MCF10A and U87 exhibit the largest parent-to-daughter displacements, reaching approximately 27 µm and 25 µm on average, respectively. In contrast, HEK293T and MDA-MB-231 display smaller displacements, averaging approximately 16 µm and 13 µm. These trends are consistent with the motility patterns observed in the displacement analysis, where MCF10A and U87 generally exhibit.

Figure 7(d) reports the average displacement of cells between consecutive observations. The trends closely mirror those observed for parent-to-daughter displacement following mitosis. Cell lines that exhibit larger inter-frame movements also tend to show larger parent-to-daughter separations at increasing gap factors. This relationship is expected because, under sparse temporal sampling, the first observable position of a daughter cell may occur long after the division event itself. As a result, the measured parent-to-daughter distance reflects not only the spatial separation generated by mitosis but also the subsequent migration of the daughter cells.

These results highlight an important challenge of low-frame-rate cell tracking. As the temporal gap between observations increases, mitosis events become increasingly difficult to identify and reconstruct because daughter cells can be observed at locations substantially removed from where the division originally occurred. Consequently, successful lineage reconstruction will require methods that account for both cell division and cell movement over extended time intervals.

### 4.2 Cell behavior under unhealthy imaging conditions

Beyond the main benchmark, we also include a smaller set of sequences acquired under unhealthy conditions. Although these sequences represent a limited portion of the dataset, we release them because they capture challenging and biologically distinct scenarios that are useful for testing the generalization of deep learning models and stress-testing existing methods.

In these sequences, cells are imaged under experimental stress. U87 does not currently include any unhealthy sequences and is therefore not discussed in this subsection.

Figure 8 shows the main properties of these cells lines regarding cell density, cumulative mitosis, and average displacement across gaps.

**Figure 8:**
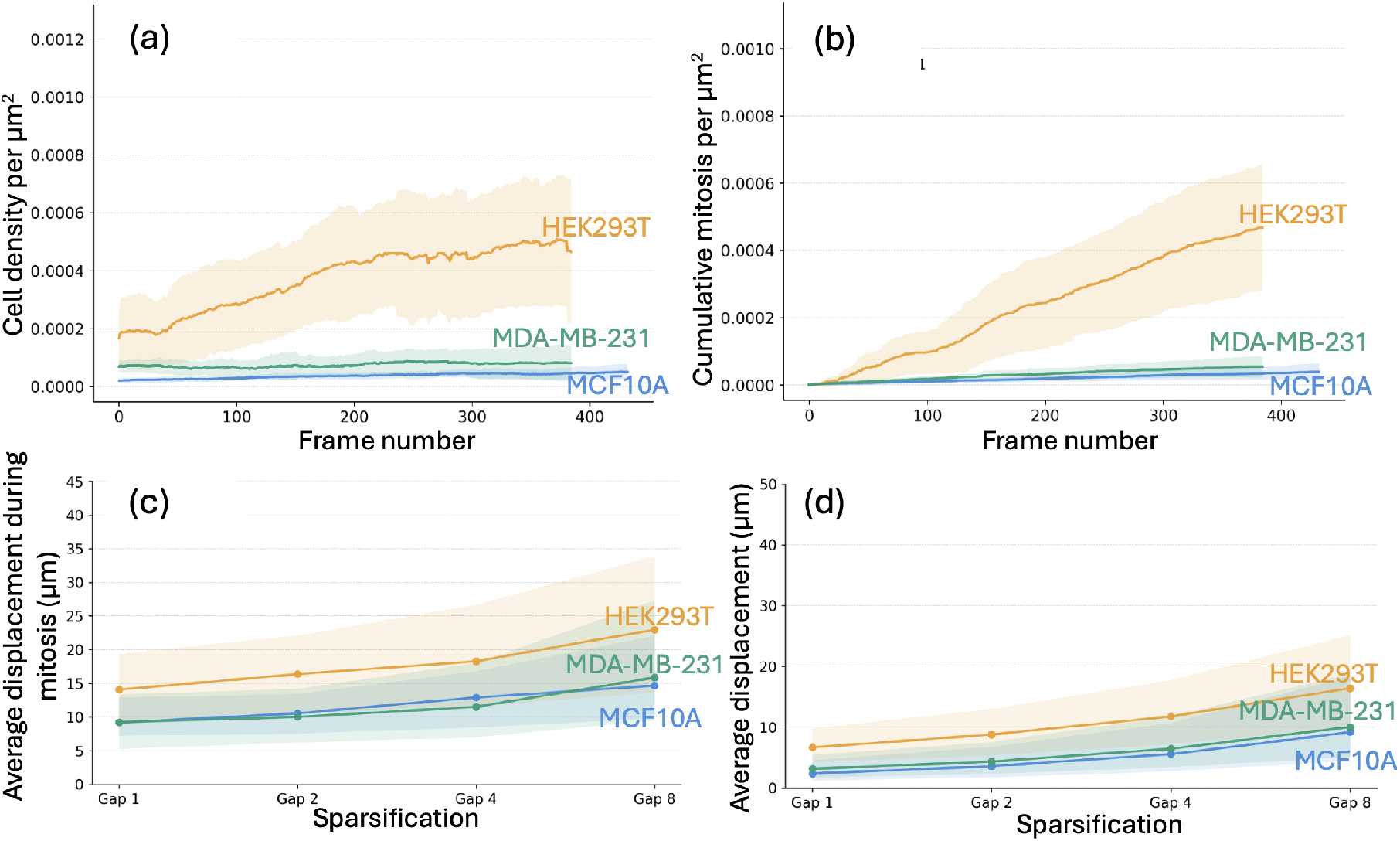
Overview of the unhealthy sequences. (a) Average cell density per unit area and (b) cumulative mitotic events per unit area measured throughout the acquisition period for each cell line. (c) Average parent-to-daughter displacement after mitosis and (d) average displacement between consecutive observations at increasing gap factors. Shaded regions represent variability across sequences of the same cell line.

The unhealthy condition affects the confluency of each cell line differently (Figure 8(a)). HEK293T still reaches the highest density, climbing from about 20 to 56 cells on average. However, this remains far below the growth observed in healthy conditions. MDA-MB-231 and MCF10A stay nearly flat in comparison, averaging only 25 to 31 cells and 10 to 24 cells per sequence, respectively.

Also mitotic actitivy is drammatically reduced across the cell lines (Figure 8(b)). HEK293T remains the most active, with the average sequence reaching 56 cumulative divisions. MDA-MB-231 reaches 21 average divisions, while MCF10A reaches 19 average divisions.

Together, these reductions confirm that the unhealthy condition suppresses both division and population expansion across the three cell lines.

As it can be expected also cell motility is affected after mitosis (Figure 8(c)) or during normal migration (Figure 8(d)). Under these conditions, HEK293T displays the highest motility, with an average frame-to-frame displacement of approximately 7.6 µm. MDA-MB-231 follows at 5.1 µm, while MCF10A is the least motile at 4.2 µm. The same ordering holds for parent-to-daughter displacement during mitosis, where HEK293T exhibits the largest separations across all gap factors and MCF10A the smallest.

Although these sequences are not included in the primary benchmark, they provide an important complementary resource for future method development. The unhealthy sequences represent a distinct regime in which cells display altered dynamics and reduced biological activity. These conditions may serve as valuable stress tests for future low-frame-rate tracking methods, enabling researchers to assess whether models generalize across different cellular states or whether their performance is tied to specific motility and proliferation patterns. Ultimately, the inclusion of these sequences may help support the development of tracking algorithms that are robust to a broader range of biological conditions and experimental settings.

## 5 Code Availability

We provide Python code and example data at https://github.com/DaVisLab/LFCT-pipeline. The repository includes functions to convert the ground-truth TrackMate XML annotations into a more human-readable CSV format; generate lineage-tree figures from the annotations; sparsify the annotations to simulate different annotation-sparsification regimes (GAP); and aggregate the corresponding phase-contrast and fluorescence frames into videos for each sparsification setting. An example sequence is also provided to demonstrate the full pipeline.

## Author contributions

M.G. performed the annotations, developed the code and lead manuscript writing. T.B. acquired imaging data and contributed to manuscript writing. X.L. acquired imaging data. C.M.G., C.K.H., and K.R.S. contributed to data analysis. M.R.B. and F.I. secured funding and contributed to manuscript writing. All authors reviewed and approved the final manuscript.

## Acknowledgments

The authors gratefully acknowledge the support provided by the National Institute Of General Medical Sciences of the National Institutes of Health under Award Number R21GM150066 and R35GM141891. Results were generated using the Palmetto Cluster at Clemson University, supported by the National Science Foundation under Grant numbers MRI 1228312, II NEW 1405767, MRI 1725573, and MRI 2018069. Any opinions, findings, and conclusions or recommendations expressed in this material are those of the author(s) and do not necessarily reflect the views of the funding agencies.

